# Objective, Quantitative, Data-Driven Assessment of Chemical Probes

**DOI:** 10.1101/168369

**Authors:** Albert A. Antolin, Joe E. Tym, Angeliki Komianou, Ian Collins, Paul Workman, Bissan Al-Lazikani

**Author notes:** Correspondence (P.W.), (B.A.-L.).

## Abstract

Chemical probes are essential tools for understanding biological systems and for target validation, yet selecting tools for biomedical research is largely biased and subjective. Here we describe the Probe Miner: Chemical Probes Objective Assessment resource – capitalising on the plethora of public medicinal chemistry data to empower quantitative, objective, Big Data-driven assessment of chemical probes. We assess >1.8m compounds for their suitability as chemical tools against 2,220 human targets and dissect their biases and limitations.

## INTRODUCTION

Small-molecule chemical probes are important tools for exploring biological mechanisms and play a key role in target validation (1–5). However, selection of chemical probes is largely subjective and prone to historical and commercial biases (2,6). Despite many publications discussing the properties of chemical probes and the proposal of ‘fitness factors’ to be considered when assessing chemical tools, scientists often select probes through web-based searchers or previous literature that are heavily biased towards older and often flawed probes or use vendor catalogues that do not discriminate between probes (2,6).

The Chemical Probes Portal (6) (www.chemprobes.org) was recently launched as a public, non-profit, expert-driven chemical probe recommendation platform and this emerging resource is already contributing to improved chemical probe selection (5). However, expert curation, by definition, can be limited in its coverage and would benefit from a complementary, rapidly updated, systematic, Big Data-driven, objective and comprehensive approach that enables researchers to keep track of the rapid advances in chemical biology-relevant data at a scale difficult to reach with expert curation – allowing comparison of the quality of large numbers of probes. Recently, a scoring system to prioritise chemical tools for phenotypic screening based on expert weighting of public and highly-curated private databases was described (7). However, a public resource that democratises Big Data-driven chemical probe assessment is still missing and would greatly contribute to target validation and mechanistic studies performed outside industry. Here, we analyse at scale the scope and quality of published bioactive molecules and uncover large biases and limitations of chemical tools in public databases. We provide the online Probe Miner: Chemical Probes Objective Assessment resource where we integrate large-scale public data to enable objective, quantitative and systematic assessment of chemical probes.

## RESULTS

### Probing the ‘Liganded’ Proteome Using Public Databases

An ambitious early grand challenge of chemical biology was to identify a chemical tool for each human gene (2,8). To assess the level of progress towards meeting this challenge, we first define the set of 20,171 curated, validated human proteins in Uniprot (9). We then utilise the canSAR knowledgebase integration (10) (http://cansar.icr.ac.uk) of major, curated, public medicinal chemistry data (including ChEMBL and BindingDB, see Methods) to determine the fraction of these proteins that are known to interact with small molecule compounds (9,10). Interestingly, we find that only 11% (2,220 proteins) of the human proteome has been ‘liganded’ (Figure 1A). This percentage is still very low even if we compare it to the 22-40% of the proteome that is estimated to be potentially druggable (Figure 1A) (10–12).

**Figure 1.**
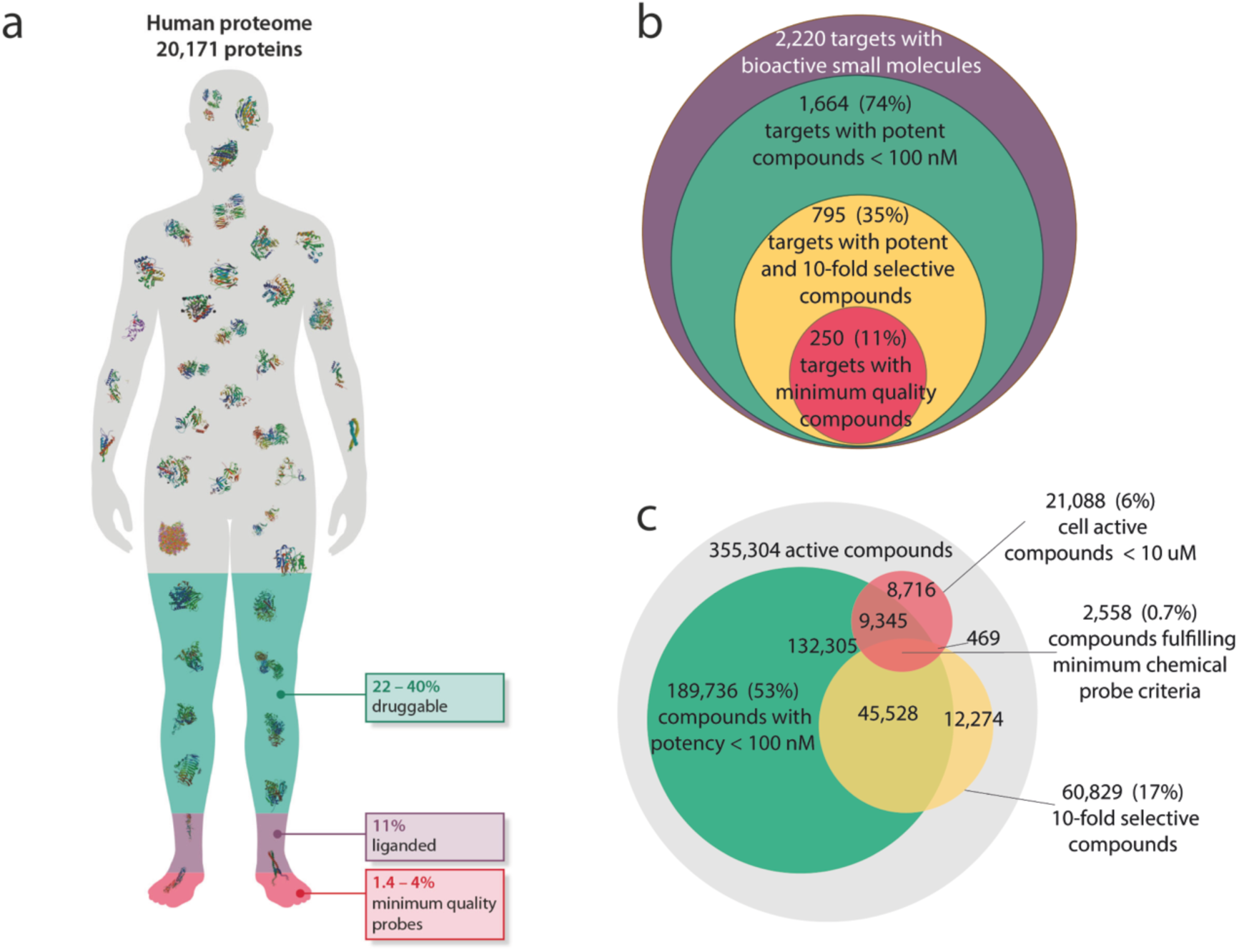
Global analysis of chemical probes as described in public databases uncovers major limitations and biases. (A) Infographic showing a human silhouette representing the human proteome and areas representing the proportion of the proteome estimated to be druggable but unliganded (green)(10,12); the proportion found to have been already liganded (purple, see Methods); and the proportion that can be studied with chemical tools fulfilling minimum requirements of potency, selectivity and permeability (red, see Methods). (B) Venn diagram illustrating the proportion of the 2,220 liganded human protein targets that can be studied with chemical tools fulfilling minimum requirements of potency, selectivity and permeability. (C) Venn diagram illustrating the number of chemical compounds fulfilling minimum requirements of potency, selectivity and permeability.

To be effective tools for mechanistic biological experiments and target validation, chemical probes must satisfy at least some basic criteria for the key properties (or fitness factors) of potency, selectivity and permeability (2). To assess how many of the compounds available in public databases would be useful in this context we establish key minimal criteria to satisfy: 1) Potency: 100 nM or better on-target biochemical activity or binding potency; 2) Selectivity: at least 10-fold selectivity against other tested targets; and 3) Permeability: As no large-scale experimental measures of permeability are available, we use cellular activity (where accessible) as a proxy and set a minimum requirement of 10 μM (see Methods). It is important to stress that these three minimal requirement levels do not guarantee that a chemical tool would be suitable for biological investigation but all suitable tools should in principle meet these basic requirements.

From the >1.8 million total compounds (TC) available in public databases, we find that only 355,305 human active compounds (HAC) have some acceptable level of biochemical activity (<10 μM) reported against a human protein. Of these, 189,736 (10.5% TC, 53% HAC) have measured biochemical activity or binding potency of 100nM or better. However, when considering selectivity, we find that only 93,930 compounds have been reported as tested against two or more targets. Of these, only 48,086 (2.7% TC, 14% HAC) satisfy our minimal potency and selectivity criteria (Figure 1C). Thus, exploration of compound selectivity in the medicinal chemistry literature appears alarmingly limited (Figure S1). Moreover, we find that the compounds that satisfy our minimal potency and selectivity criteria allow the research community to probe only 795 human proteins (4% of the human proteome) and at best 18% of the estimated druggable proteome (Figures 1A-C). Finally, when also considering cellular potency of 10μM or better, we find that the number of minimal quality probes is reduced even further to 2,558 (0.7% HAC). Under these combined criteria, compounds fulfilling minimum requirements would allow the research community to probe with real confidence only 250 human proteins (Figure 1B). This accounts for an unacceptably low percentage (1.2%) of the human proteome.

The amount of information available for a target will clearly impact any statistical analysis of its corresponding chemical tools. To assess the role of differing levels of experimental characterisation, we define the ‘Information content’: For each target, A, we collect all small molecules (C) shown to be active against this target. For each compound we then count the number of additional targets (T) against which it has been tested, regardless of activity level. Thus, the information content is IC_A_=ΣCT. (See Methods for detail). As expected, we find large biases in the amount of data in public medicinal chemistry databases available for different targets. We also observe a wide range in the number of compounds fulfilling our minimum criteria among targets (0-204, Figure S2). For example, some targets have many well-characterised compounds, several of which fulfil our minimum criteria; e.g. the metalloprotease ADAM17 that has 1,586 active compounds of which 31 satisfy our minimal criteria. Other targets have large numbers of compounds with differing degrees of characterisation, yet few, if any, satisfy our minimal criteria; e.g. CDK5 has 1,849 active compounds, none of which satisfy our minimal criteria with the data available (Figures S2 and S3). Several factors could influence the observed biases, for example the availability of selective probes varies significantly across the analysed targets (0-896 selective compounds). The identification of selective probes may be simpler for some targets that have distinctive binding sites (e.g. PPARγ) and difficult for others that share binding sites with numerous family members (e.g. ABL1). Increasing the number of large-scale panel screens of many compounds against many targets will certainly help complete the information matrix required to identify good quality probes. Indeed, half of the 50 targets with the greatest number of minimum quality probes are kinases, which benefit from systematic library screens (Figure S2). However, this brute-force approach alone is insufficient. Overall, we find poor correlation (R^2^ = 0.1) between the number of reported experimental measurements and the number of minimum quality probes (Figure S3), indicating that our community needs to be smarter in designing and testing compounds in addition to increasing the throughput of data generation.

### Probing Disease Genes

Our systematic approach allows us to investigate, more globally, how well existing chemical tools equip us to probe mechanistically the function of disease genes – which is particularly important for therapeutic target validation. As an exemplar, we analyse data for a set of 188 cancer driver genes (CDG) with activating genetic alterations (13) and examine the availability of minimal quality chemical probes for these drivers. We find that 73 (39% CDG) have already been liganded, and of these 25 (13% CDG) have chemical tools in public databases fulfilling minimum requirements of potency, selectivity and permeability (Table S1, Figure S4). Since these genes include a large number of well-studied disease genes, these results represent a substantial improvement over the pattern in the whole proteome (Figure 1B) but nevertheless 87% of CDG do not have a minimum quality chemical tool (Table S1). Moreover, the vast majority of tools concentrate on few targets, further demonstrating the documented trend to focus research efforts in areas of science already well-studied (Table S1, Figure S4) (14). This analysis further uncovers a severe lack of chemical probe availability and significant bias where tools are available.

### Objective Assessment of Chemical Probes

Given biases and limitations discussed above, it is imperative that researchers can comprehensively access all the data publicly available in an objective and data-driven manner in order to select the best characterised chemical probes available for their target of interest and to understand their liabilities and limitations at the start. To this end, we describe a scoring metric that utilises >3.9m bioactivity data points publicly available in canSAR (10) to rationally prioritise chemical probes.

To create a metric that allows objective, data-driven ranking of all compounds tested for a particular target, we developed a set of six scores mirroring our previously described ‘fitness factors’ (2). Namely, Potency Score, Selectivity Score, Cell Score, Structure-Activity Relationship (SAR) Score, Inactive Analog Score and PAINS Score (see Methods, Figure S5). We predefine a default weighting of these scores – which emphasises the importance of potency and selectivity (see Methods). However, we also provide the facility for researchers to customise and adapt the weights to suit their own questions and expertise.

Using our default scoring scheme allows us to highlight compounds that make good candidates for chemical probes, and defines their key limitations. For example, we assess 1,694 compounds for PIK3CB (Figure S6). The five top ranking probes include the clinical candidate pictilisib or GDC-0941 (top rank) (15) and the frequently used probe PI-103 (2^nd^ ranked) (16) (Figure 2, Figure S6), both widely profiled in large kinase panels. However, our scores highlight that both these compounds have certain selectivity liabilities due to cross-PI3K activity (Figure 2, Figure S6). The PIK3CB/PIK3CD inhibitor AZD-6482 ranks 10^th^ (Figure 2), due to its partial PI3K selectivity towards PIK3CA and PIK3G (17) and most other PIK3CB-selective chemical series are also represented among the top scoring probes (17).

**Figure 2.**
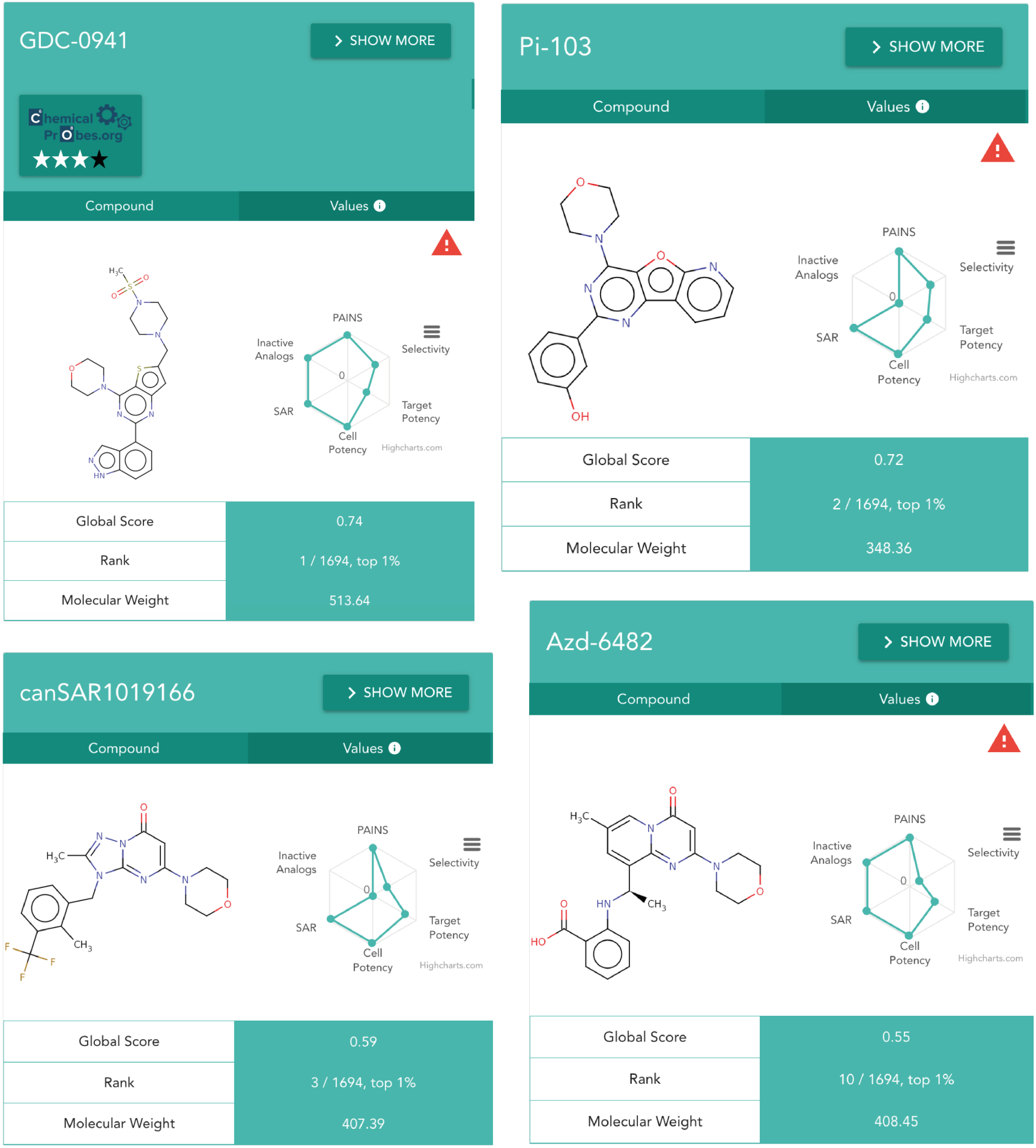
Top ranked PIK3CB chemical probe cards comprising the chemical structure and the radar plot with the corresponding chemical probe scores. Probes also curated by The Chemical Probes Portal include their expert ratings in a 4-star format and, when compound is not 10-fold selective against another protein, a danger icon appears to alert the researcher that there might be selectivity liabilities when using those compounds as PIK3B chemical probes.

Furthermore, our systematic assessment additionally highlights recently published compounds such as canSAR1019166 (18) which ranks 3^rd^ using our default scoring (Figure 2, Figure S6). This compound is both potent and, unlike pictilisib and PI-103, is selective against other PI3K proteins. Since no reports of screening canSAR1019166 against wider kinase panels are in the public domain yet, other selectivity liabilities may emerge in future. Additionally, this compound may not be commercially available. There are also key compounds whose biochemical characterisation is not provided in public medicinal chemistry databases, and thus it is not possible to appropriately assess them using our approach. For example, this is the case for the PIK3CB-selective clinical candidate GSK-2636771 (17,19), which is currently not top ranked in our resource.

### Probe Miner: A Public Resource For Objective Assessment of Probes

To empower the community to utilise this objective approach, we developed the regularly and automatically updated Probe Miner: Chemical Probes Objective Assessment website (http://probeminer.icr.ac.uk; Figure 3). This is a user-friendly, interactive web-based resource that allows access to the data and probe rankings, as well as full customisation of the scores and the ability to deep-dive into the data.

**Figure 3.**
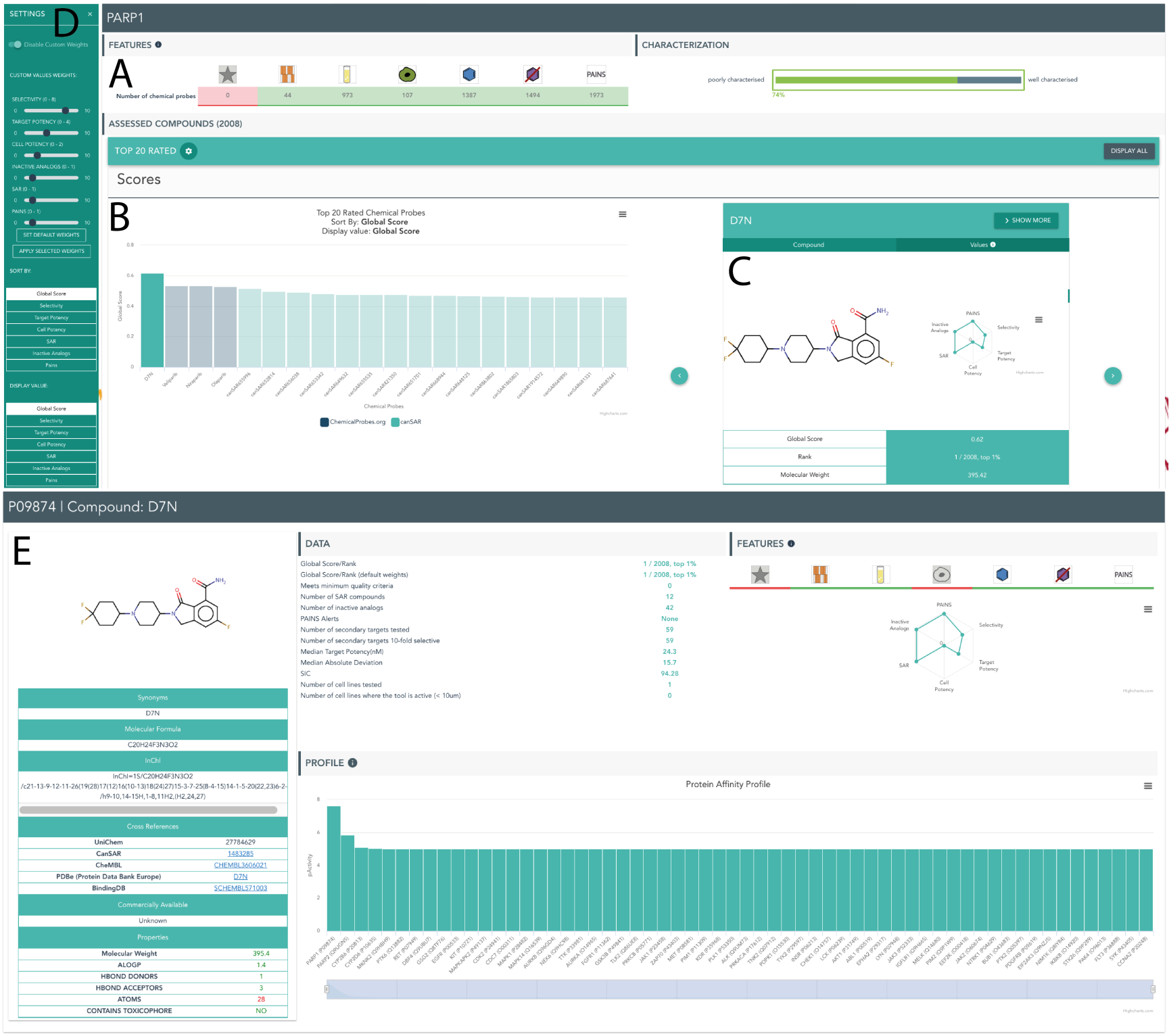
Probe Miner resource. Snapshot of the overview and chemical tool pages of the resource using the human target PARP1 as an example. (A) summaries of the data and statistical analyses using our algorithm. Coloured icons provide immediate visual indication of the overall quality and liabilities of compounds for this target. (B) Easy-to-navigate distribution of the twenty top ranking probes. (C) A compound viewer interactively linked to the distribution which shows the chemical structure and key information for the probe, as well as the values of the six score components as a radar plot. (D) Easy-to navigate settings panel to enable customisation of Global Score, displays and rankings. (E) Individual chemical probe pages where detailed information is provided including links to other resources, commercial availability, raw data to generate the scores and a target profile to provide an overview of compound selectivity (further details in Figures S7-S11).

Probe Miner is a target-centric, systematic probe assessment resource. We provide an interactive graphical overview page for each target (Figure 3, Figure S8) which comprises three major sections: A) summaries of the data and statistical analyses using our algorithm. Coloured icons provide immediate visual indication of the overall quality and liabilities of compounds for this target. B) Easy-to-navigate distribution of the twenty top ranking probes as well as tools to customise the scores, weights and ordering of probes. C) A compound viewer interactively linked to the distribution which shows the chemical structure and key information for the probe, as well as the values of the six score components as a radar plot. As Probe Miner is intended to complement the Chemical Probes Portal, we highlight the expert-curated probes that are also assessed in our resource and provide links to their individual pages in the Chemical Probes Portal. Figure 3 shows the large-screen version of the resource. The website adapts to multiple devises and screen sizes.

It is important to stress that the selection of the ‘best’ probes must be always tailored to the scientific question under investigation and, therefore, the final decision on which tools to use must always be undertaken by the researcher. Therefore, as introduced above, we enable researchers to set the weights of each of the individual fitness factor scores that contribute to the Global Score (Figure 3D and Figure S11), so that these can be adapted to the requirements of user needs. Through our easy-to navigate settings panel (Figure 3D), we also enable customisation of displays and rankings. Nonetheless, for convenience, we also provide our pre-defined weighting for the Global Score (See Methods).

From this overview page, researchers can navigate to individual probe pages where detailed information is provided (Figure 3E, Figure S9). All the raw data necessary to generate the scores is accessible in a tabular format, together with the radar plot displaying the scores, the ranking and further detailed information. The target profile is also provided as a bar plot displaying the median affinities for the given compound against all the targets it has been screened against to enable a quick and easy understanding of the selectivity of the compound for the target of interest. Finally, cross-references to key public resources are also provided, including canSAR, ChEMBL, BINDINGDb, The Protein Databank, and The Chemical Probes Portal as well as synonyms and commercial availability where available (Figure 3E, Figure S9). Finally, a Chemical Probes Table page enables access to all the chemical tools and raw data for each target in a tabular format, their filtering, and full download of all the data to enable the chemical biology community to develop these scores further (Figure S10). Overall, this resource provides a public online framework for objective, Big-Data driven prioritization of chemical tools. We will maintain the resource and perform automatic updates following the release of new versions of the public databases that are integrated.

The power of our resource is the objective, systematic, regularly updated assessment that relies on public medicinal chemistry databases. However, as illustrated throughout our analysis, limitations in data availability or curation can pose a significant challenge in some cases. It is our belief that arming researchers with the information and highlighting potential areas of error or bias is key to empowering them in making the best decisions. For example, to alert researchers to cases where selectivity may be a problem, we have incorporated a danger icon that warns researchers when a chemical probe fails the 10-fold selectivity criterion against another protein. The target profile of every compound can also be easily accessed at the chemical probe page for a quick visual understanding of the actual selectivity of each chemical tool for the target of interest (Figure S9) while links to The Chemical Probes Portal are provided and expert curation highlighted to draw attention on probes recommended by experts. Moreover, our objective assessment performed at scale can identify compounds that we rank as good probe candidates, but are not currently curated in The Chemical Probes Portal nor are they commercially available and highlight them to vendors for commercial provision.

To help address errors and inaccuracies in public databases, we are undergoing a continuous curation exercise of the underlying data and have established an email where we can be contacted by any researcher that identifies such errors or inaccuracies affecting the objective assessment of chemical probes. Overall, even high-quality public databases are not exempt from errors and inaccuracies that are extremely challenging to identify and fix. Data-driven approaches rely on the quality of the data they use and it is thus paramount that we as a community address the errors and inaccuracies of public databases to maximize the benefit from this costly generated data.

### Benchmarking against The Chemical Probes Portal

Using our predefined Global Score, we compare the top ranked chemical probes with the expert-curated probes available in the Chemical Probes Portal (6). For this analysis, we focus on the selective probes curated by The Chemical Probes Portal and assigned to no more than two targets within the Portal (see Methods). From the 133 probes rated by the Chemical Probes Portal (date: 06/02/2017; see Methods), 71 are associated with no more than 2 targets and recommended by experts (Rating ≥ 3, see Methods). Of these 71 probes, 46 curated probes, corresponding to 45 targets, can be mapped to public databases. Thirty-one (67%) of the 46 probes rank among the top 20 probes by our predefined Global Score and 18 (39%) rank among the top 5 (Table S2). Our analysis of the 15 expert-recommended probes that fail to reach the top ranking uncovers that the incompleteness of data available in public databases (often because the probe was published in a non-indexed journal) and also inaccuracy of public data are the major limitations (Tables S2 and S3). As the purpose of our resource is to compliment the Chemical Probes Portal with strictly objective data-driven information, we explicitly exclude any curated probes that have no data in the underlying medicinal chemistry knowledgebases. However, to address the broader lack of coverage of chemical biology data in public medicinal chemistry databases, the canSAR knowledgebase is actively expanding to curate key missing literature. In future this growing knowledgebase will enhance our objective assessment and increase the overlap between our resource and The Chemical Probes Portal.

Importantly, our analysis highlights 193 compounds with high rankings (in the top 5) that are not yet curated within the Chemical Probes Portal, and that may complement the tools available to explore these targets. This highlights the clear synergies of combining the large-scale objective assessment of all available compounds with indepth expert curation. As a result of this synergy, we are now collaborating with the Chemical Probes Portal to share information and for example to recommend probes identified by our objective assessment method for expert curation at the Portal (see example below).

### Use Cases: PARP-1, CHEK2 and OPRK1

The Poly(ADP-ribose) polymerase 1, PARP1, exemplifies the complementarity of the two resources and illustrates how they can be used by researchers to empower the selection of the best probes. From the five pan-PARP probes recommended by the Chemical Probes Portal, olaparib, veliparib and niraparib are highly ranked by our predefined Global Score (Figure 3 and 4). Unfortunately, key information is missing in public resources regarding the probes AZ0108 and E7449, the latter published in a journal not indexed in public databases (20). Accordingly, these two probes are not highly ranked in our resource (Figure 4). On the other hand, our objective assessment identifies another probe that scores at the top but has not yet been curated by the Chemical Probes Portal. NMS-P118 is a recently published PARP1-selective inhibitor that was also comprehensively screened for kinase selectivity following reports of off-target kinase activity among PARP inhibitor (21,22). Therefore, NMS-P118 emerges as a potential candidate with which to probe specifically for PARP1 (Figure 4). Based on our findings, we proposed NMS-P118 to The Chemical Probes Portal and this chemical probe is now under review for expert curation. The example of PARP inhibitors illustrates the utility and complementarity of our objective approach at prioritizing new probes that could later on be assessed by chemical biology experts using wider information and also uncovers the need to better capture the chemical biology literature in public databases.

**Figure 4.**
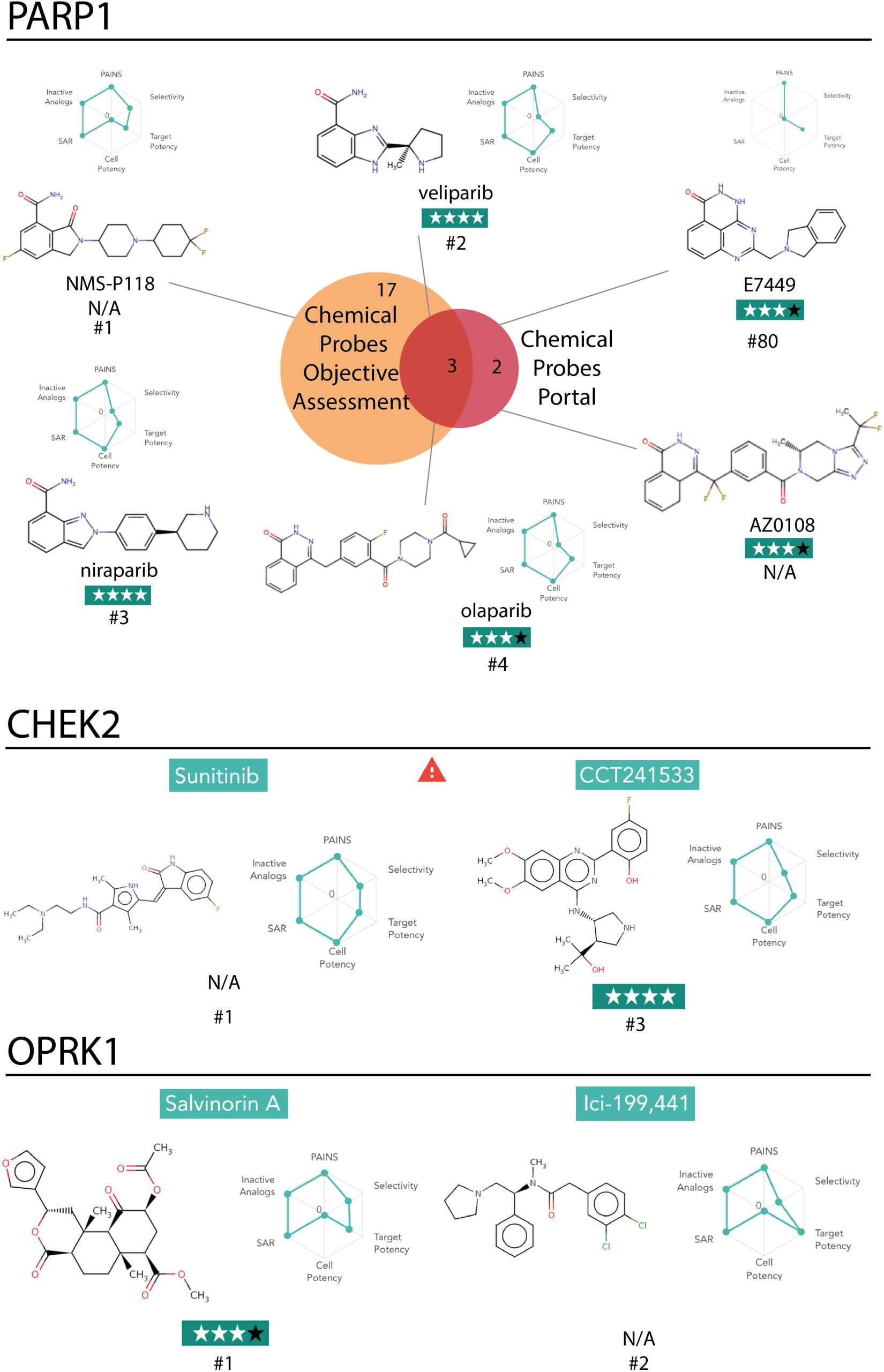
Analysis of the ranking of chemical probes for the targets PARP1, CHEK2 and OPRK1. On top, Venn diagram comparing the PARP1 chemical probes recommended by the Chemical Probes Portal (see Methods) and the Probe Miner resource as ranked by the pre-defined Global Score. Chemical structures are displayed, as well as names, radar plot showing the six Chemical Probe Scores, Chemical Probes Portal reviewers’ rating and Probe Miner ranking when available. On the bottom, top-ranked probes for CHEK2 and OPRK1.

The serine/threonine-protein kinase CHEK2, exemplifies cases that uncover the limitations in the public medicinal chemistry data resources. For CHEK2, our algorithm failed to prioritise the highly selective chemical probe CCT241533 (23) while ranking as the first chemical tool the promiscuous but highly characterised kinase inhibitor sunitinib. When investigating the reasons behind this ranking, we found that the affinity of CCT241533 for CHEK2 had been wrongly curated in ChEMBL, making the probe appear non-selective (Figure S12). We reported this error to ChEMBL and it has now been corrected in both ChEMBL and our canSAR database. Now, CCT241533 ranks as the third best probe in Probe Miner (Figure 4). It is important to stress that our default weighting rewards probes that have been extensively characterised over those that have not. For example, if one probe shows selectivity but has only been tested against two off-targets, while another probe is not completely selective but has been tested against hundreds of targets, our selectivity score – that balances the amount of selectivity information with the actual selectivity – may be higher for the widely characterised probe, depending on the number of targets tested and the selectivity units (see Methods for details).

This is the case in our analysis of sunitinib as a probe. It has been very thoroughly characterised across the kinome results in this compound ranking as the top probe when using our pre-defined Global Score, as in this case the Selectivity Score gives a very high score to this probe (see Methods and Figure S13-S17). This emphasises the difficulty of comparing chemical tools that have been screened against very different number of targets (or when this information is differently covered in public databases) and the importance that researchers are aware of these differences to make informed choices. Our resource highlights these issues and alerts the user to them as described above.

The use of more than one chemical probe with different chemotypes is strongly recommended for mechanistic studies and target validation (5). Accordingly, our third example the Kappa opioid receptor, OPRK1, illustrates how Probe Miner can be used to identify second probes and potentially contribute to increase the completeness of The Chemical Probes Portal. For OPRK1 there is only one chemical probe covered in The Chemical Probes Portal, Salvinorin A. While Probe Miner also identifies Salvinorin A as the top-ranking probe for OPRK1, it also identifies chemically distinct probes for this target such as ICI-199,441 (24) that is potent, selective and commercially available and can thus be used in conjunction with Salvinorin A to probe for the OPRK1 receptor (Figure 4). We recommend the OPRK1 chemical probe ICI-199,441 for evaluation by The Chemical Probes Portal for expert review that complements our data-driven and objective assessment while increasing the coverage of OPRK1 inhibitors by The Chemical Probes Portal.

These and other examples demonstrate that Probe Miner, when used in conjunction with The Chemical Probes Portal, enables the researcher to make informed choices in the selection of probes for their experiments.

## DISCUSSION

Through our systematic analysis of >3.9 million experimental activities for >1.8million compounds in curated public medicinal chemistry databases, we provide objective, data-driven systematic scoring of 355,305 compounds against 2,220 human targets. Using our systematic, objective, Big Data-driven assessment, we demonstrate that the majority of human proteins lack minimal quality small-molecule chemical tools that are needed to probe their function. This number can only improve over time with increased availability of experimental data. As such, to empower researchers to select the best tools available to them at any time, a regularly updated, objective ranking derived from the evolving underlying medicinal chemistry data is key. This systematic view, combined with expert-curated knowledge from the Chemical Probes Portal, provides both the breadth and depth required to make informed choices on which chemical probes to select.

Our Probe Miner resource represents, to our knowledge, the first publicly available resource enabling objective, data-driven and systematic assessment of chemical tools. Providing regularly updated analyses of all compounds in public medicinal chemistry databases, it democratises access to this knowledge and enables the wider community to make informed choices using up-to-date information. We demonstrate that our objective prioritization of chemical tools aligns largely with expert recommendation. Moreover, our approach also synergises with expert curation in that it analyses hundreds of thousands of compounds on a regular basis and objectively highlights probes with promise and for further curation by experts.

While the available public data provides an excellent starting point for such objective analysis, we also uncover incompleteness and inaccuracies of data deposited in public databases that limit the full benefit this approach. The public databases used in this analysis were developed mainly for medicinal chemistry applications and, accordingly, many chemical biology publications are not covered. Moreover, we have identified an error in public databases that has now been corrected (Figure S12) and also found several inaccuracies, particularly regarding the annotation of cell-based EC_50_s as biochemical IC_50_s (Table S3). Therefore, there is a great need to better capture and curate chemical biology information from the literature in public knowledgebases if we are to make the most of these expensively generated data.

Here, we analyse large-scale public data to catalogue and examine the quality of currently available chemical tools and facilitate their data-driven and objective assessment. We use this approach to rank and highlight many promising compounds against >2000 human targets. We find major limitations in the description and characterisation of chemical tools in public databases, highlighting our limited knowledge of chemical tool selectivity and the large biases identified. It is therefore paramount that the chemical biology community both improves the characterisation of currently available chemical tools and also continues the development of new tools to study areas of biology that remain poorly characterized in order to address the historical limitations and biases. In this context, to empower researchers in the selection of chemical tools and compliment expert-recommendation approaches, we have created and describe here the Probe Miner resource as the first public, live, data-driven, systematic, objective, chemical probe assessment engine.

This resource can be used with a predefined weighting of fitness factors – of which measures of biochemical activity/binding potency and selectivity, as well as surrogate measures of cell permeability, are given greater weighting in the overall score. The present weightings for assessing and ranking chemical probes may be especially useful for users such as bioscientists who are not chemical biology or medicinal chemistry experts – which is a group identified as requiring advice and user-friendly resources to help in selecting chemical tools for exploring biology and target validation (5).

Alternatively, the weighting can be customised according to individual research needs. For example, expert users may wish to alter the weighting of fitness factors to suit their own experience and opinion. Or they may wish to vary the weightings of different factors to see how this affects the ranking of probes.

Despite the limitations in data accuracy and completeness that we and others have identified in public databases (7), we demonstrate that objective data-driven prioritization generally aligns well with expert recommendation at The Chemical Probes Portal when the information is accurate and available in public databases. We view it as a major strength that Probe Miner, which provides value through its automatic updating capability and its breadth across the whole liganded proteome, complements the expert view provided by The Chemical Probes Portal.

The Chemical Probes Portal currently has about 400 probes and aims to grow this as quickly as possible. In addition to use in own right, our new Probe Miner resource can be used by the community to fill the gap while the Portal expands into protein families that have not yet been covered, and can also be employed to help prioritize probes for subsequent expert curation and assessment at the Portal or by individual researchers. In the longer term we propose that the Probe Miner and the Chemical Probes Portal have complementary strengths which will make their continued use synergistic and mutually beneficial to the user community.

In conclusion, we demonstrate here that objective, quantitative, Big Data-Driven large-scale assessment based on public data can contribute to improving overall evaluation and prioritization of chemical probes. This will help empower researchers towards much-needed improvement in the selection of chemical tools for biomedical research and target validation.

## AUTHOR CONTRIBUTIONS

A.A.A., I.C., P.W. and B.A.-L. designed the research. A.A.A. performed the experimental work. A.A.A., I.C., P.W. and B.A.-L. conducted data analysis and interpretation. A.A.A., J.E.T., A.K. and B.A.-L. designed the website. J.E.T and A.K. coded and developed the website resource. A.A.A., I.C., P.W. and B.A.-L. contributed to manuscript preparation.

## ACKNOWLEDGMENTS

The work was primarily supported through the People Programme (Marie Curie Actions) of the 7th Framework Programme of the European Union (FP7/2007-2013) under REA grant agreement no. 600388 (TECNIOspring programme); the Agency of Business Competitiveness of the Government of Catalonia, ACCIO to A.A.A.; and the Wellcome Trust Sir Henry Wellcome Postdoctoral Fellowship (204735/Z/16/Z) to A.A.A.; The work utilises the canSAR Knowledgebase which is funded by a CRUK Strategic Project Award (C35696 / A23187) to B.A.-L. In addition, B.A.-L., I.C and P.W are funded by The Institute of Cancer Research (ICR) as well as The Cancer Research UK (CRUK) grant to the Cancer Research UK Cancer Therapeutics Unit (grant C309/A11566). P.W is a CRUK Life Fellow. The authors thank many of their collaborators for discussions and valuable input into the preparation of this manuscript. In particular, the authors thank Amy Donner for helpful discussion with regard to the Chemical Probes Portal.

## METHODS

### 1. Definitions

During this work we have used the following definitions:

#### 1.1 Target

Human protein that is known to interact with a chemical compound.

#### 1.2 Reference Target

Since a chemical compound can bind to multiple protein targets and we score each compound-target pair, the reference target is defined as the target of the compound that is being evaluated.

#### 1.3 Potency Score

Score that measures the potency of the biochemical interaction between each compound – target pair.

#### 1.4 Selectivity Score

Score that measures the selectivity of each compound – target pair. Selectivity is one of the most important properties that a chemical tool should fulfil in order to be useful to study the biological function and therapeutic potential of a specific protein (2,4). However, it is challenging to measure due to large biases in the number of targets screened per each molecule (Figure S13) and thus the selectivity score balances our actual knowledge of selectivity with the amount of selective information available.

#### 1.5 Cell Score

Since no large-scale experimental measure of permeability is available, we use cellular activity as a proxy. Accordingly, the Cell Score is a binary score that measures whether a given chemical molecule is known to be active in cells, and thus accounts not only for the permeability but also for the solubility and cell activity fitness factors because when a compound is active in a cell line assay we assume it fulfils minimum requirements of permeability and solubility to modulate the target of interest in cells (2).

#### 1.6 SAR Score

Structure-Activity Relationships (SAR) increase confidence that the biological effect of a given chemical tool is achieved via the modulation of the reference target. Accordingly, the SAR score is a binary score measuring whether there are (SAR Score = 1) known SAR for the compound - reference target pair (2).

#### 1.7 Inactive Analog Score

Inactive analogs can be useful controls to rule out off-target effects. Therefore, the Inactive Analog Score is a binary score measuring whether there are known inactive analogs for the compound – reference target pair (2).

#### 1.8 PAINS Score

Pan-assay interference compounds (PAINS) are compounds that interfere with the detection methods of screening assays and are thus problematic artefacts that have been identified to be widely used in many scientific publications as chemical tools, thus leading to the wrong conclusions.(25) In an effort to contribute to discourage the use of these compounds in chemical biology, the PAINS score measures whether a compound is predicted to be PAINS-free (PAINS Score = 1). However, given the recent alerts on false-positives among compounds predicted to be PAINS using current methods (26), it is important to stress that this score is a prediction and thus should be taken with caution. Accordingly, we enable researchers to set the weight of the PAINS Score in the Global Score according to their expert judgement and the specific assay that they are using.

### 2. Chemical Probe Scores & Measures

#### 2.1 Potency Score

We consider bioactivity data on compound-target pairs integrated in the knowledgebase canSAR (that integrates high quality bioactivity data from CHEMBL and BINDINGDB) (27,28) where the target type is a protein, the protein is human as defined by its associated Uniprot ID and the units can be transformed to ‘nM’ (9,10). We calculate the median of all the reported values for each compound-target pair distinguishing between ‘=’ and ‘>’ values and transform them into pActivity values (- log(Activity[Molar])). We consider active all compounds with a median pActivity below 10,000 nM. Compounds with conflicting ‘=’ and ‘>’ data for the same target are considered inactive. Given that only 0.7% of bioactivities are below 100 picomolar but they strongly bias the normalisation of the potency score, these activity values (pActivity ≥ 10) are given a value of 10. For all the compound-protein pairs considered as active, the potency score is calculated as the normalization of the pActivity values in a scale from 0 (pActivity = 5) to 1 (pActivity ≥ 10). There have been arguments against any numerical aggregation of potency values due to their wide variation across biological systems and technologies, but the proposed alternative uses subjective expert weighting of several databases including highly curated proprietary databases that are not widely accessible (7). We have thoroughly investigated cases where a wide distribution of potency values has been reported, such as nilotinib (CHEMBL255863) (7). We identify that these wide distributions are often due to inaccuracies in the annotation of cellular ECs0s as biochemical IC_50_s and the annotation of data from mutants as wild type proteins (27). We have calculated the Median Absolute Deviation (MAD) of the potency median calculated for each compound-target pair and we identify that these large variations affect only a very small number of compounds (3.6%; Figure S14). We provide the MAD to facilitate the identification of cases where this wide distribution may affect the reported performance of the chemical tool. These results support the use of high quality public databases for the assessment of chemical tools and highlight the need to better curate these high quality databases to make the most of this expensively generated data.

#### 2.2 Selectivity Score

For each compound, we calculate a different selectivity score for each of the proteins it interacts with (median pActivity > 5) considering all of their compound-target interactions. In order to balance the actual knowledge of selectivity with the amount of information available (Figure S13), the Selectivity Score is composed by three different factors in an attempt to reflect our limited cataloguing of selectivity following the formula below:

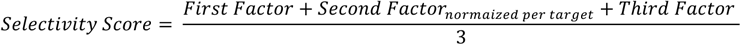

The First Factor accounts for the actual knowledge of selectivity. In order to calculate the First Factor, the number of off-targets that the compound has been screened against is calculated first, without including the target being evaluated. Second, the median pActivity values are used to discern whether there is at least 10fold selectivity (1 log unit) between the potency of the compound for the reference target and the potency for each off-target (pActivity_ReferenceTarget_ – pActivity_off-Target_ ≥ 1). The 10-fold selectivity cut-off has been previously used as the minimum selectivity requirement to consider that a chemical probe is selective (29). If there is 10-fold selectivity between the potency of the compound for the reference target and the potency of the off-target, this off-target is considered a selective off-target (Figure S15). Next, we calculate the number of selective off-targets. The First Factor of the Selectivity Score is obtained by dividing the number of selective off-targets by the total number of off-targets, using the following equation:

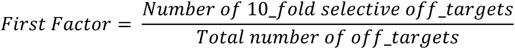

Therefore, if a compound-reference target interaction is at least 10-fold selective against any other target, the value of the first factor will be 1. In contrast, if the compound-target interaction of interest is not 10-fold selective against any other compound-target interactions, the first factor will be lower than 1 (Figure S15). If there is no information regarding any off-target, the Selectivity Score is set to 0.

The second factor of the score is a measure of the amount information available regarding selectivity. This second factor distinguishes between compounds that have an equal first factor but have been screened against a (very) different number of targets. Moreover, it also balances the actual knowledge of selectivity - measured by the first factor - with the amount of information available regarding selectivity that can be very different between different compounds, challenging their comparison (Figures S16 and S17). In order to calculate it, we have developed a measure of selective information that we have termed Selectivity Information Content (SIC). The SIC is calculated as the summary of the differences between the median pActivity of the reference target and the pActivity of each off-target minus one:

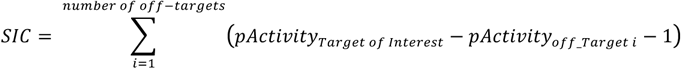

This strategy enables the evaluation of the selectivity information from the 10-fold selectivity cut-off in order to distinguish selective from un-selective information that will be positive and negative, respectively (Figure S16). Therefore, in the final summary unselective data compensates for selective data. Interestingly, the SIC could also be regarded as the number of selectivity units from the given compound-protein interaction. To calculate the Second Factor of the Selectivity Score, the SIC is divided by the number of targets that would be modulated at the same time since there is no sufficient selectivity between them, therefore the number of unselective off-targets plus the target of interest, following this formula:

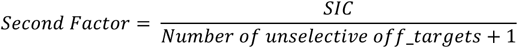

This division enables to reduce the SIC of compound-target interactions that are very high, as the compound has been screened against a very large number of off-targets, but represent suboptimal compounds to probe for the target of interest as they inhibit several other targets with similar or higher affinity (Figure S17).

The second factor is finally normalized within each target as we observe that different targets can have very different SIC ranges. The main reason for this is that there are target families such as kinases where familywide profiling is very common while this is not common for other target families and a global normalization would profoundly bias the results.

Finally, the last factor measures the percentage of the proteome that the compound has been screened against, which is generally very low, with the following formula:

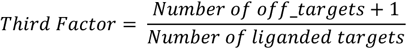

Therefore, this factor works as a reminder that the selectivity of chemical tools is generally a big unknown. Ultimately, the three factors are added and the final score is normalized (Equation 1). Overall, the Selectivity Score is able to balance different flavours of selectivity but how compounds screened against a very different number of targets should be prioritized remains a philosophical question. Therefore, our aim at developing the selectivity score was to prioritize compounds and facilitate the evaluation of the information available but the final decision should always be taken by the chemical biologist after careful evaluation of available information and tailored to the requirements of the specific experiment.

#### 2.3 Cell Score

To calculate the Cell Score we compute the median of all compound – cell line bioactivities reported in canSAR that can be transformed to ‘nM’ (10). We consider a compound has positive Cell Score (Score =1) if it is active in at least one cell line considering a cut-off of 10,000 nM (median pActivity > 5). This cut-off is set to minimise the risk of considering non-specific drug toxicity that may lead to cell death at high concentrations. Compounds that have activity values less potent that the cut-off or that have not been tested in cell line assays are given a Cell Score value of 0.

#### 2.4 SAR Score

To calculate the SAR Score we first calculate the level 1 of the scaffold tree for all compounds in canSAR as it has been described to have advantages over other scaffold definitions (30). Next, we consider a compound – reference target pair has SAR (SAR Score = 1) if there is at least another compound reported in the same publication (identical PubMedID) with the same level 1 scaffold active against the reference target (pActivity > 5).

#### 2.5 Inactive Analog Score

The Inactive Analog Score measures whether there are compounds sharing the level 1 scaffold with the compound being evaluated that are reported to be inactive (pActivity < 5) for the reference target.

#### 2.6 PAINS Score

We apply PAINS rules to filter compounds that are given a PAINS alert by giving them a PAINS score of 0 (25).

#### 2.7 Global Chemical Probe Score

The Global Chemical Probe Score is a combination of the previous 6 Chemical Probe Scores with customizable weights to allow chemical biologists to prioritize the best chemical tools for the specific requirements of their experiments. We have predefined weights for a case where selectivity is twice as important as potency, which in turn is twice as important as cell activity, which in turn is twice as important as SAR, inactive analogs and PAINS scores. However, we want to stress that different proposed experimental cases will require different weights of these scores in order to access the best probes. We do not think that there is a unique Global Score applicable to all chemical biology experiments and accordingly the weights to each of the scores can be personalised for individual user needs in the website resource. Note that it is unfortunately not possible to fairly compare our Global Score to the recently developed TS score for prioritisation of chemical tools for phenotypic screening as TS uses expert weighting of several databases, including highly-curated proprietary databases for which we do not have access (7). The Global Score has the following formula (for pre-defined weights a = 8, b = 4, c = 2, d = e = f = 1):

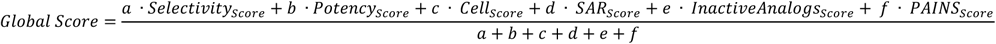

#### 2.8 Commercial Availability

Commercial availability is not reported as a score because we believe that this would discourage the supply of the best chemical tools and does not represent an inherent property of compounds. However, we recognise that knowing whether a chemical tool is commercially available is important for chemical tool selection and thus we provide this information in each chemical tool synopsis page. We consider a compound is commercially available if it is present in the catalogue of eMolecules (https://www.emolecules.com/) that comprises over 8 million compounds from a network of suppliers. To identify if a compound is present in the eMolecules database we use UniChem cross-references (31).

### 3 Development of the Probe Miner: Chemical Probes Objective Assessment Resource

We have developed an open website (http://probeminer.icr.ac.uk) using PHP, HTML and jQuery JavaScript library to enable public access to the Probe Miner resource as a framework for chemical probe prioritization using data integrated from publicly available knowledgebases.

#### 3.1 Target Icons

To facilitate a rapid and intuitive evaluation of chemical probe quality we have adapted the chemical probe scores to a binary representation and developed a set of icons that can be shown in colour or in grey scale depending on the chemical tool fulfilling certain criteria. These icons are displayed in each chemical tool synopsis page (Figure S9). Moreover, in order to facilitate a target's-eye view of chemical tool quality using these icons, the number of probes fulfilling these criteria is also displayed below the icons in each Target Overview page (Figure S8a). A description of each icon can be found in the following table:

| Icon Name | Description | Image |
| --- | --- | --- |
| Target Selectivity | Exemplified by an (orange) heatmap icon, it shows whether a compound inhibiting this protein is screened against at least one other target and has at least 10-fold selectivity against any other target. |  |
| Target Potency | Exemplified by a (yellow) test tube icon, it shows whether a compound inhibits this target with at least 100 nM potency. |  |
| Cell Potency | Exemplified by a (green) cell, it shows whether a compound binding to the target of interest is active in a cell line with at least 10 μM potency. | |
| Minimum Standard | Exemplified by a (red) star, it is an aggregate of the three previous scores (that themselves are independent from each other), measuring whether there are compounds inhibiting this target with minimum standards of target potency ($p\text{Activity} \geq 7$), selectivity (at least one tested off-target and 10-fold selectivity against any off-target) and cell potency (activity below 10 μM in at least one cell line) simultaneously. It is a key icon showing whether a compound fulfilling these minimum quality requirements is found in publicly available databases. | |
| SAR | Exemplified by a (blue) benzene ring, it measures whether there is at least one compound binding to this target that has SAR as defined by the SAR Score (vide supra). |  |
| Inactive Analog | Exemplified by an (indigo) barred benzene ring, it measures whether there is at least one inactive analog of the compound as defined by the Inactive Analog Score (vide supra). |  |
| PAINS | Exemplified by a (violet) 'PAINS' icon, it shows whether there is a compound inhibiting this target that has no PAINS alerts as defined in the PAINS Score. |  |

#### 3.2 Target Information Content Score

In order to inform on the amount of information available, we develop a measure of the information content for each target, not only in terms of the number of chemical compounds screened against it but also their characterization in terms of selectivity. Accordingly, for each target, every compound screened against it is counted as one unit of information. Moreover, for each compound tested against that target, each other target the compound was screened against is also counted as another unit of information. Therefore, each target has a final information value that accounts for the number of screened compounds plus the number of other targets each compound was screened against.

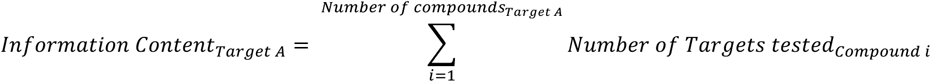

Next, each target is ranked according to their information value. The information content score reports the percentile of each target in terms of ranking, being 100% for the targets with the highest information values and 0% for the targets with the lowest information values.

### 4 Analysis of the ‘liganded’ proteome and chemical tools for cancer genes

The Potency Score (vide supra) is used to calculate how many human proteins interact with a chemical molecule with a median activity below 10,000 nM (pActivity < 5) and thus represent the currently liganded proteome. The Potency Score, Cell Score, the number of off-targets and the number of selective off-targets calculated for the Selectivity Score are subsequently used to calculate how many compound – target interactions fulfilled minimum chemical probe requirements. Only compound - target interactions with median pActivity ≤ 7, reported to have an affinity below 10,000 nM in at least one cell line, screened against at least one other target and at least 10-fold selective against all other targets screened are selected. Finally, the absolute number of human protein targets and chemical molecules selected is calculated. In order to compare information content with chemical tool quality, information values calculated for the Target Information Content Score (vide supra) are compared to the number of compounds fulfilling minimum chemical probe requirements for each target (Figure S3). For the analysis of minimum-quality chemical tools for cancer driver genes we extracted the chemical tools fulfilling minimum requirements from the previous analysis and annotated to the 188 cancer targets identified as potentially driving cancer in a recent pan-cancer analysis (13).

### 5 Analysis of chemical probes from the Chemical Probes Portal

All the chemical probes from the Chemical Probes Portal are downloaded from the Chemical Probes Portal website (http://www.chemicalprobes.org/browse_probes; downloaded 06/02/2017) including key information such as name, target(s) names, PubChem CID and Average Recommendation (Table S2) (6). Probes are mapped to canSAR compound IDs when possible using the provided PubChem CIDs, ChEMBLIDs or SMILES. The most potent target from the reported values is considered for the analysis and mapped to UniprotIDs via the provided gene names (Table S2). The oldest Primary Reference of the probe is also recorded and mapped to PubMed ID, Journal name and publication year (Table S2). Since our assessment is performed at the single target level, we focus on the selective probes for the comparative analysis. From the 133 probes available in the resource at the accession date, 109 were associated to no more than two targets (Table S2). Next, we selected probes that have a SAB Rating ≥ 3 and are thus recommended by experts following the Chemical Probes Portal guidelines (http://www.chemicalprobes.org/sab-rating-system). From the 109 selective probes, 71 are recommended by experts. From the 71 recommended probes, 46 could be mapped to publicly available medicinal chemistry databases and have affinity data for the primary target that enables the calculation of the scores. It is worth noting that many of the probes that could not be mapped were published in 2016 or 2017 and they had not yet been included in public databases such as ChEMBL. From these 46 probes, their ranking for their intended target is calculated according to our predefined Global Score (Table S2). In 30 cases (65%), the recommended probes are ranked among the top 20 by the predefined Global Score and in 17 cases (37%) the recommended probes rank among the top 5. The analysis of the 15 probes that are recommended by The Chemical Probes Portal but do not rank among the top 20 by the Global Score uncovers that the main reasons for not ranking correctly are data incompleteness (mainly because key information was published in a non-indexed publication) or data inaccuracy (mainly EC_50_s curated as IC_50_s) (Tables S2 and S3).

